# Podoplanin interaction with caveolin-1 promotes tumour cell migration and invasion

**DOI:** 10.1101/488304

**Authors:** Luisa Pedro, Jacqueline D. Shields

## Abstract

Podoplanin, a highly O-glycosylated type-1 transmembrane glycoprotein, found in lymphatic endothelial cells, podocytes, alveolar epithelial cells and lymph node fibroblasts is also expressed by tumour cells, and is correlated with more aggressive disease. Despite numerous studies documenting podoplanin expression, the mechanisms underlying its tumour-promoting functions remain unclear. Using a murine melanoma cell line that endogenously expresses podoplanin, we demonstrate interactions with proteins necessary for cytoskeleton reorganization, adhesion and matrix degradation, and endocytosis/receptor recycling but also identify a novel interaction with caveolin-1. We generated a panel of podoplanin and caveolin-1 variants to determine the molecular interactions and functional consequences of these interactions. Complementary *in vitro* and *in vivo* systems confirmed the existence of a functional cooperation in which surface expression of both full length, signalling competent podoplanin and caveolin-1 are necessary to induce directional migration and invasion, which is executed via PAK1 and ERK1 pathways. Our findings establish that podoplanin signalling mediates the invasive properties of melanoma cells in a caveolin-1 dependent manner.

**Summary Statement:** This manuscript describes a new interaction and functional cooperation between podoplanin and caveolin1 that drives tumour cell invasion into surrounding tissues.

## Introduction

Podoplanin is a small, highly glycosylated mucin-like protein, expressed in a range of tissues, and is now associated with diverse range of functions depending upon tissue type; in podocytes podoplanin is associated with glomerular filtration barrier integrity, while in lymphatics it is critical for development and lymph sac budding, where it’s disruption prevents segregation from the cardinal vein and lymphovascular defects (Herzog et al., 2013; Uhrin et al., 2010). In mature lymphatics, associated with CD44, podoplanin can promote endothelial migration, and in lymph node FRCs it interacts with CLEC2-expressing dendritic cells supporting their movement within lymph nodes (Acton et al., 2012). Its expression may be induced by TGFβ (Suzuki et al., 2008), and signals downstream of PMA, RAS, and Src stimulation (Nose et al., 1990; Shen et al., 2010). Further to chemical cues, podoplanin up-regulation has been reported in osteoblasts in response to mechanical cues from mineralization and tissue stiffening (Prideaux et al., 2012).

Podoplanin is emerging as a marker of pathologies including arthritis (Takakubo et al., 2016) and cancer. Up-regulation has been reported in a growing number of cancers including melanoma (Ochoa-Alvarez et al., 2012), lung, colorectal, oral (Martin-Villar et al., 2005) and breast (Wicki et al., 2006) cancers, where it is frequently associated with poor prognosis. Thus, there is increasing interest to understand how it exerts pro-tumour functions, and whether it represents a viable therapeutic target. Studies using genetically modified cells have correlated podoplanin with enhanced metastatic capacity via platelet aggregation (Kunita et al., 2007; Takagi et al., 2013), promotion of EMT (Martin-Villar et al., 2015), or destabilization of cell-cell junctions in the absence of EMT (Wicki et al., 2006). Expression of podoplanin is often restricted to the leading edge, where it is in contact with surrounding tissue and remodelled extracellular matrix, implying that podoplanin up-regulation may occur in response to environmental cues from neighbouring stroma and promoting invasion.

Antibodies targeting the extracellular domain of podoplanin reduced lung colonization (Kaneko et al., 2012; Kato et al., 2006); possibly by blocking signalling functions or interfering with interactions between podoplanin and other surface proteins. In light of this, we sought to investigate the mechanisms by which podoplanin can drive invasion using cells with endogenous expression, identifying new binding partners underlying for this behaviour. Although having no impact on tumour proliferation or survival, we identified that podoplanin induced significantly increased directional motility and invasive capacity of melanoma cells via activation of PAK1. These activities required functional signalling capacity within the cytoplasmic tail of podoplanin and functional co-operation with caveolin-1 at the cell surface lipid rafts.

## Results

### Podoplanin expressing tumour cells are more invasive *in vivo*

To determine the impact of podoplanin upon the invasive potential of tumour cells we utilized B16.F10 melanoma cells that endogenously express low levels of podoplanin (Fig. 1a). We generated variants to either knockdown or overexpress podoplanin (Fig. 1a-b), which translated to varying degrees of surface expression (Fig. 1b). Mice with intravenously administered tumour cells consistently developed a greater number of lung nodules (Fig. 1c-d) and tumour burden (Fig. 1e) with increasing levels of surface podoplanin. However, in contrast to the metastatic setting, levels of podoplanin on tumour cells at the primary site had no impact on growth (Fig. 1f), indicating that podoplanin was not essential for tumour growth but has the capacity to impact downstream events.

**Figure 1:**
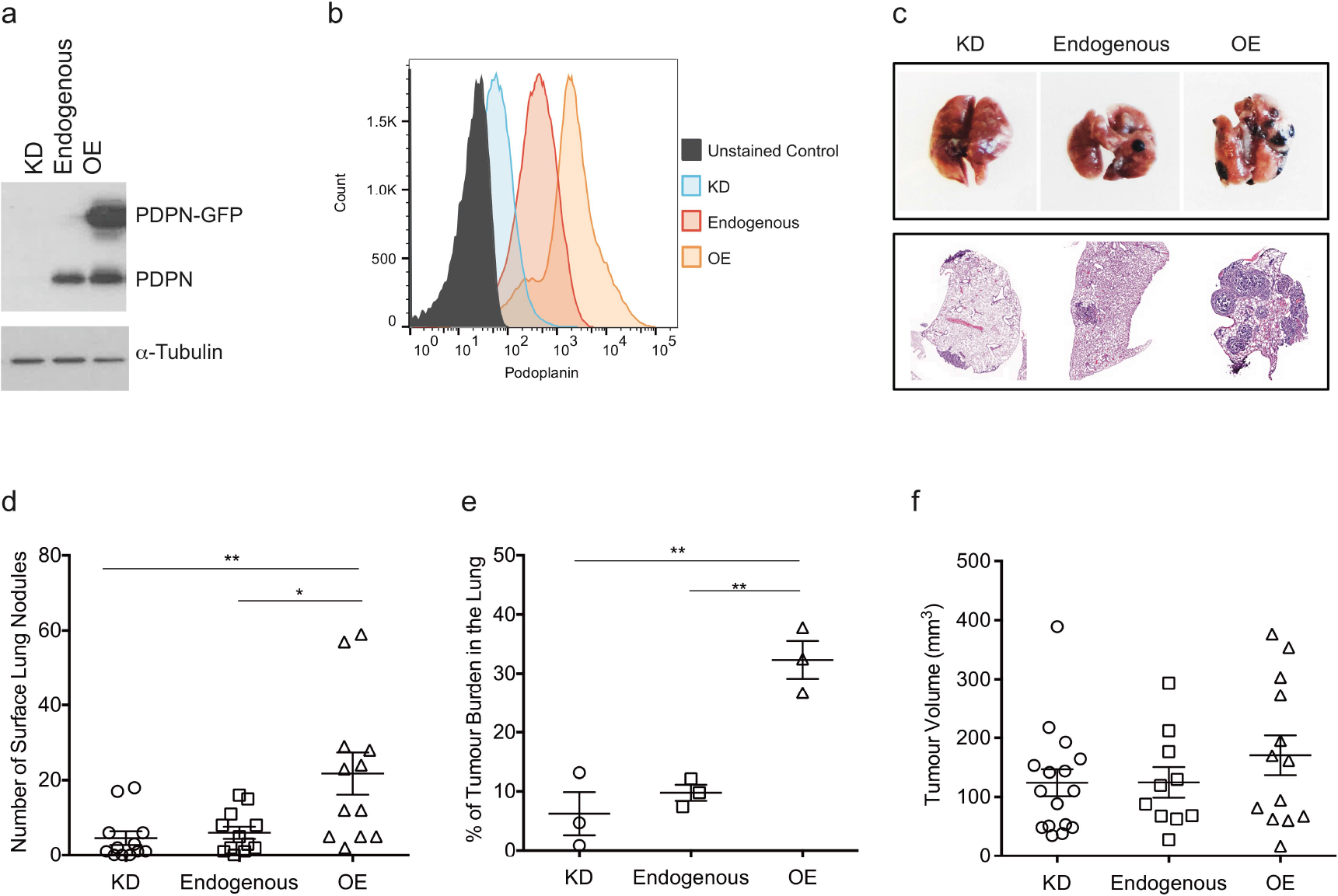
Podoplanin levels correlate with lung metastatic burden but not with primary tumour size. a) WB analysis showing total protein levels of the three different B16.F10 cell lines with different podoplanin levels: KD (knockdown), no podoplanin expression; Endogenous (low), endogenous expression of podoplanin; OE (overexpression), high level of podoplanin expression. b) Representative flow cytometry histogram illustrating levels of surface podoplanin present on the different cell lines. c) Representative images of lungs from experimental metastasis assays. Lung nodules present 21 days after intravenous B16.F10 injection (top) and haematoxylin/eosin stain images of lung sections illustrating tumour burden (bottom). d) Quantification of the number of tumour nodules per lung. n= 4 independent experiments performed in triplicate. e) Representative data for quantification of H&E sections for percentage of tumour burden. f) Volume of primary tumours 9 days after subcutaneous implantation of B16.F10 cells with different podoplanin levels. n= 3 independent experiments performed in triplicate. Data presented as Mean ± SEM. *p<0.05, **p<0.01 (One-way ANOVA with Bonferroni post hoc test).

### Podoplanin expressing tumour cells are more invasive *in vitro*

To identify the mechanisms underlying these functional differences, we investigated consequences of podoplanin levels on tumour cell behaviour *in vitro*. No impact on tumour cell proliferative capacity (Fig. 2a and Fig. S1a), viability (Fig. S1b-c) or cell motility, as assessed by random migration on a 2D surface (Fig. 2b), was observed with varying podoplanin expression. In contrast, when a stimulus, in this case a wound was applied, the amount of surface podoplanin supported significantly increased wound closure velocity in a dose dependent manner (Fig. 2c). Similarly, enhanced invasion through trans-wells with thick collagen gels (data not shown) and of 3D spheroids into surrounding matrix were observed with increasing podoplanin (Fig. 2d).

**Figure 2:**
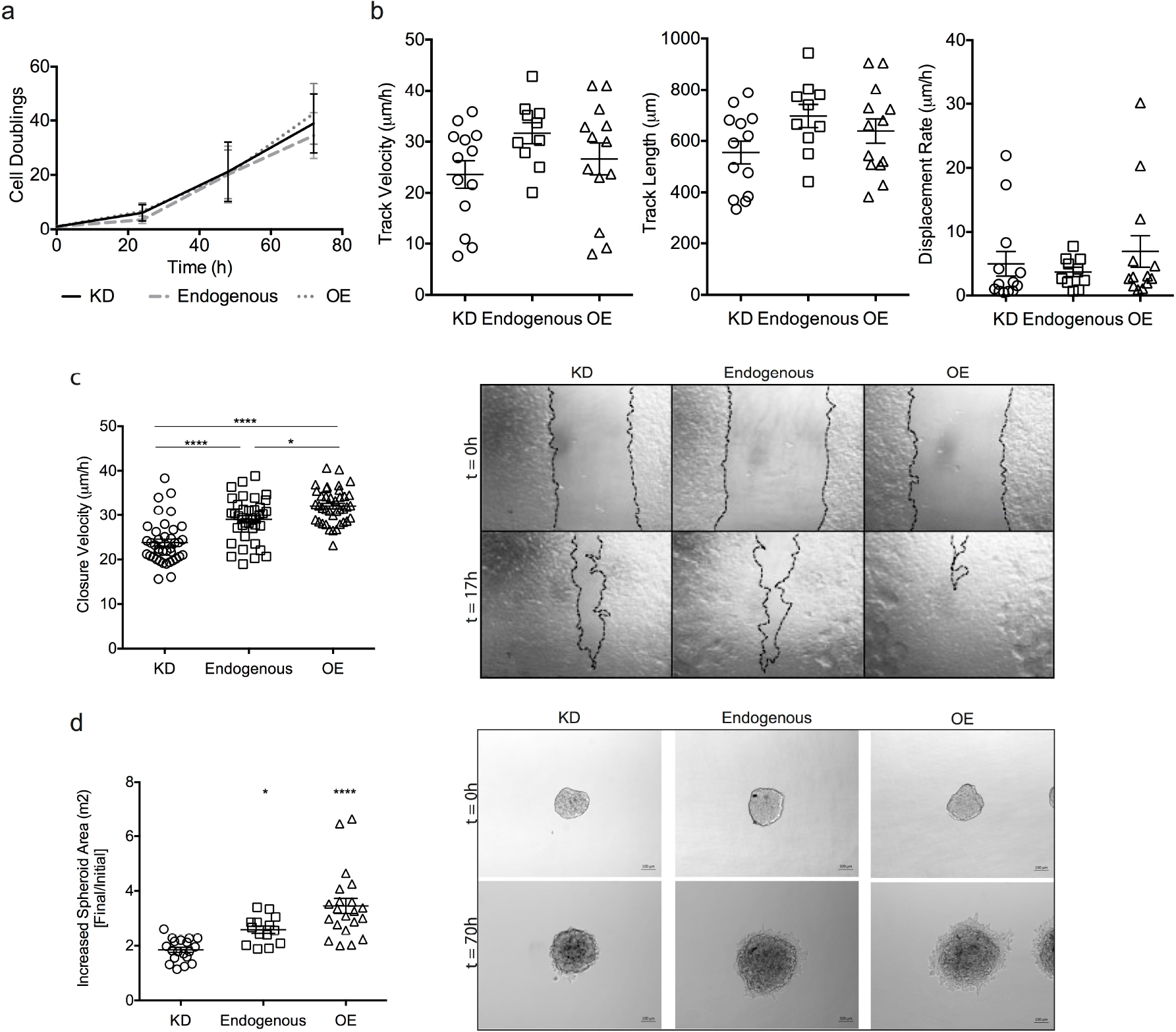
Podoplanin expression supports directional migration *in vitro*. a) Growth curve for the different podoplanin-expressing cells, represented as cell doubling over 72 hours. n=3 independent experiments performed in triplicate. b) Quantification of random migration of B16.F10 variants. Data are represented as track velocity (μm/h), track length (μm) and displacement rate (μm/h). Representative data for 3 independent experiments performed in triplicate. Each point represents a cell. c) Quantification of directional migration in response to a wound stimulus. Data presented as scratch assay linear closure velocity of the scratch (μm/h). n=4 independent experiments performed in triplicate where measurements from 3 different positions of each well were plotted overtime. Representative snapshots from wound scratch movies at t=0 and 17h post scratch (right). d) 3D invasion assay using spheroids composed of 1000 cells seeded in a 3D matrigel matrix and followed overtime. Data presented as spheroid size increase after 70 hours in culture using Zen software (Zeiss). Representative spheroid images at t=0 and 70 hours post seeding (right). Scale bar 100μm. n=3 independent experiments performed in triplicate. Data presented as Mean ± SEM. *p<0.05, ****p<0.0001 (One-way ANOVA with Bonferroni post hoc test).

### Podoplanin signals via PAK1 to support cytoskeletal rearrangement

In addition to impaired directional velocity of tumour cells with decreasing podoplanin, siRNA illustrated that disruption to podoplanin altered the morphology of migrating cells (Fig. 3a). Cells migrating away from the leading edge no longer exhibited a mesenchymal phenotype with individual leading cells driving gap closure, instead a more rounded phenotype was observed at the collectively migrating front (Fig. 3a). As cellular migration is dependent on interactions between many different proteins, but its regulation is exerted by the activity balance of small Rho GTPases, namely RhoA, Rac1 and Cdc42, we analysed their activity in samples where a wound stimulus had been applied. Consistent with previously published data (Wicki et al., 2006), although total levels of RhoA, Rac1 and Cdc42 increased with podoplanin, their activity decreased (Fig. 3b). This led us to believe that podoplanin-mediated directional migration uses an alternative pathway. qPCR arrays for cytoskeleton regulators and motility related genes allowed for a more targeted screening of mediators of podoplanin pro-invasive functions, and identified several significantly deregulated genes following podoplanin disruption (Table 1). Of note, PAK1 and ERK1 expression and activity increased with podoplanin (Table 1 and Fig. 3c). To determine whether podoplanin-driven directional migration signals in a PAK-1-dependent manner we used kinase inhibitor 1,1′-Dithiodi-2-naphthtol (IPA-3) to prevent PAK1 signalling. In knockdown and low podoplanin-expressing (endogenous) B16.F10, inhibition of PAK1 had no significant effects on migration compared with vehicle controls (Fig. 3d). In contrast, IPA-3 dose-dependently impaired migration and wound closure in podoplanin-overexpressing tumour cells (Fig. 3d). The maximum concentration examined was 10 μM since doses beyond this affected proliferation and viability (Fig. S2a). As this inhibitor may exert some off target effects with other PAK signal transducers at higher concentrations, we then disrupted PAK1 expression with siRNA. Consistent with earlier data, neither podoplanin nor PAK1 knockdown had a significant impact on cell migration in KD cells. In overexpressing cells, however, PAK1 siRNA significantly impaired scratch closure velocity to levels equivalent with podoplanin knockdown (Fig. 3e) indicating a role for PAK1 signalling downstream of podoplanin in the induction of directional cell migration. Moreover, inhibition of PAK1 alone was enough to promote delocalization of podoplanin from the cell membrane (Fig. 3f and Fig. S2b).

**Figure 3:**
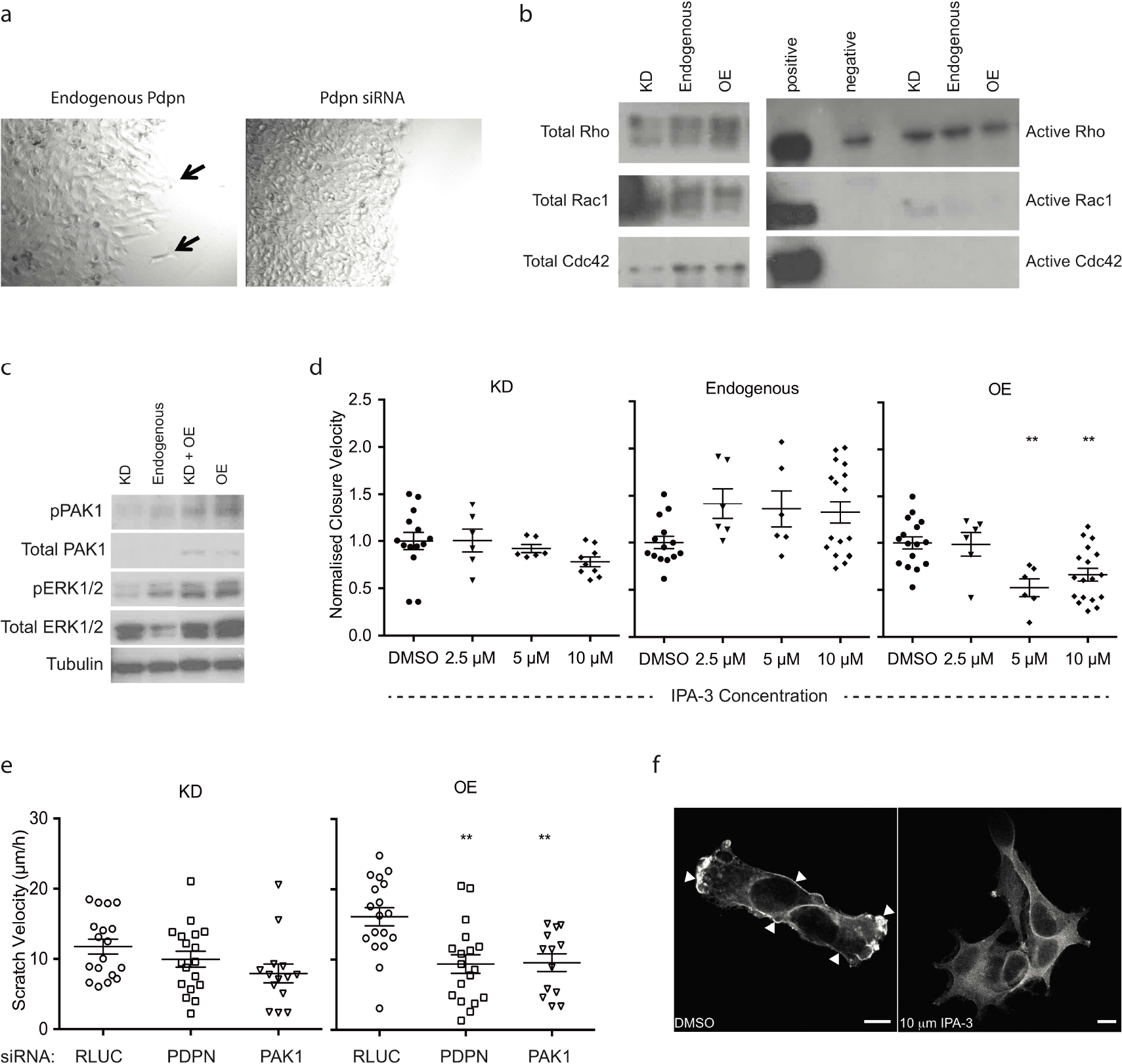
PAK1 drives cytoskeletal rearrangements in podoplanin-expressing B16.F10 cells. a) Representative images of leader cells closing a scratched wound, for cells expressing podoplanin (endogenous, arrows) or cells with silenced podoplanin expression (siRNA). b) Representative western blot data for total and active GTPases (Rho, Rac1, and Cdc42) for cells with different podoplanin expression levels. GTPase activity assays were performed at least three times. c) Representative western blot detection of pPAK1, Total PAK1, pERK1/2, Total ERK1/2, and Tubulin for cells with the different podoplanin expression levels. d) Quantification of directional migration of B16.F10 variants in response to a wound stimulus following PAK1 inhibition. Data presented as normalized linear closure velocity. n=2 independent experiments performed in triplicate. e) Directional migration of tumour cells following Pdpn and PAK1 knockdown with siRNA. Data shows linear wound closure velocity. n=3 independent experiments performed in duplicate. f) Localization of podoplanin in B16.F10 cells with and without treatment with 10μm of IPA-3. Representative confocal image. Arrows indicate podoplanin ‘hotspots’ at membrane extremities. Scale bar: 10μm. Data presented as Mean ± SEM. **p<0.01 (One-way ANOVA with Bonferroni post hoc test).

**Table 1.** Genes with a fold change greater than 1.5.

| **Cytoskeleton Regulators Array [Qiagen PAMM-088]** |  |  |  |
| --- | --- | --- | --- |
| **Gene Bank** | **Description** | **Symbol** | **Fold Change** |
| NM_011035 | P21 protein (Cdc42/Rac)-activated kinase 1 | Pak1 | 25.5844 |
| NM_009707 | Rho GTPase activating protein 6 | Arhgap6 | 1.8115 |
| NM_027180 | ArfGAP with RhoGAP domain, ankyrin repeat and PH | Arap1 | 1.7079 |
| NM_139300 | Myosin, light polypeptide kinase | Mylk | 1.7079 |
| NM_177710 | Slingshot homolog 2 (Drosophila) | Ssh2 | -1.5082 |
| NM_145575 | Caldesmon 1 | Cald1 | -1.6968 |
| NM_009515 | Wiskott-Aldrich syndrome homolog (human) | Was | -1.8123 |
| **Motility Array [Qiagen PAMM-128]** |  |  |  |
| **Gene Bank** | **Description** | **Symbol** | **Fold Change** |
| NM_007616 | Caveolin-1, caveolae protein | Cav1 | 2.8219 |
| NM_028810 | Rho family GTPase 3 | Rnd3 | 1.9908 |
| NM_134156 | Actinin, alpha 1 | Actn1 | 1.8747 |
| NM_133796 | Rho GDP dissociation inhibitor (GDI) alpha | Arhgdia | 1.8319 |
| NM_011577 | Transforming growth factor, beta 1 | Tgfb1 | 1.7013 |
| NM_007778 | Colony stimulating factor 1 (macrophage) | Csf1 | 1.632 |
| NM_007858 | Diaphanous homolog 1 (Drosophila) | Diap1 | 1.6058 |
| NM_008610 | Matrix metalloproteinase 2 | Mmp2 | 1.6058 |
| NM_011101 | Protein kinase C, alpha | Prkca | 1.5948 |
| NM_011949 | Mitogen-activated protein kinase 1 | Mapk1 | 1.5728 |
| NM_178046 | Supervillin | Svil | 1.5619 |
| NM_013599 | Matrix metalloproteinase 9 | Mmp9 | 1.5404 |
| NM_175260 | Myosin, heavy polypeptide 10, non-muscle | Myh10 | 1.5298 |
| NM_010513 | Insulin-like growth factor I receptor | Igf1r | 1.5087 |
| NM_008006 | Fibroblast growth factor 2 | Fgf2 | 1.5018 |
| NM_008404 | Integrin beta 2 | Itgb2 | -3.3326 |

### Podoplanin interacts with caveolin-1

PCR arrays revealed a significant increase in caveolin-1 mRNA with podoplanin (Table 1). Following this observation, and since caveolin-1 has also been reported at the invasive front, associated with poor prognosis and is implicated in cell migration (Felicetti et al., 2009; Yoo et al., 2003), we investigated this further. Membrane fractionation confirmed that both podoplanin and caveolin-1 co-localized within lipid rafts of B16.F10 cells (Fig. 4a). Moreover, immunoprecipitation (IP) confirmed that podoplanin and caveolin-1 physically interact (Fig. 4b). This was verified by reciprocal IP (Fig. 4c). Confocal analysis further demonstrated co-localization of caveolin-1 and podoplanin to the cell surface (Fig. 4d top panel). Interestingly, disruption of podoplanin expression by siRNA resulted in coincident reduction of caveolin-1 at the cell surface (Fig. 4d mid panel). In contrast, disruption of caveolin-1 had little effect on podoplanin localization, indicating that podoplanin may be dominant for the lipid raft interaction (Fig. 4d lower panel). Functionally, interference to either podoplanin or caveolin-1, and therefore disruption of any interaction, resulted in a significant impairment of directional migration in B16.F10 OE cells but no effect was observed for the KD cells (Fig. 4e). Moreover, overexpression of caveolin-1 was not sufficient to rescue wound closure velocity (Fig 5d). Together, these data imply that the presence of podoplanin-caveolin-1 complexes at the cell surface are required for directional tumour cell migration in response to an external stimulus which is not caveolin-driven, however disruption to either is sufficient to prevent a response.

**Figure 4:**
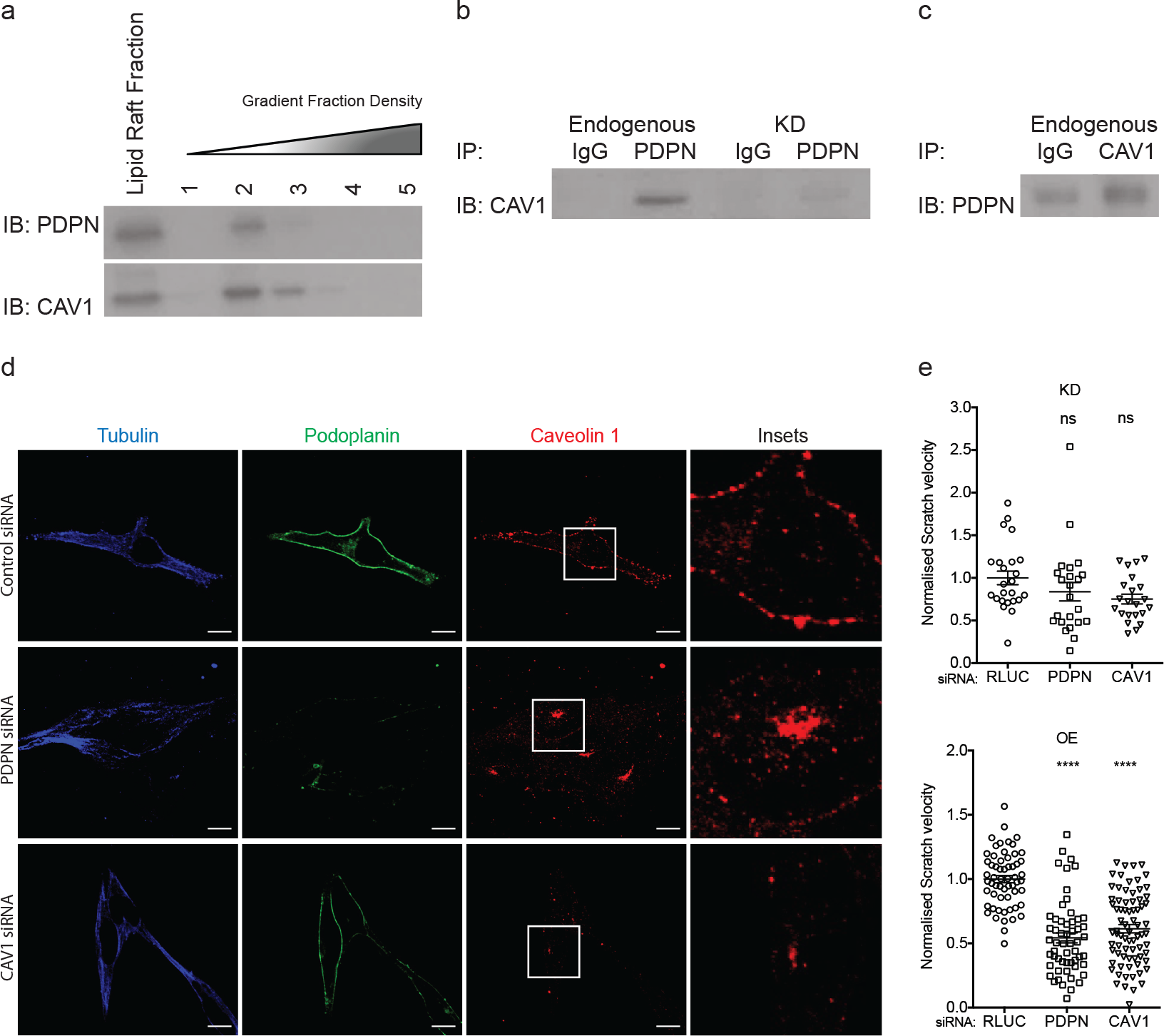
Podoplanin and caveolin interactions determine caveolin-1 localization. a) Western blot analysis of lipid rafts extracted from B16.F10 cells. Fractions were probed for podoplanin and caveolin-1. b) Immunoprecipitation of podoplanin from B16.F10 and B16.F10 KD cell extracts and probed for caveolin-1. c) Reciprocal immunoprecipitation of caveolin-1 from B16.F10 cell extracts probed for podoplanin. d) Representative confocal images stained for tubulin, podoplanin and caveolin-1 following siRNA silencing of Pdpn or Cav1. Insets: close up view of caveolin-1 cellular localization. Scale bar: 10μm. e) Quantification of directional migration of B16.F10 and B16.F10 KD after wounding. Data presented as normalized linear closure velocity of the scratch. n=4 independent experiments performed in triplicate. Data shown as mean ± SEM. ****p<0.0001 (One-way ANOVA with Bonferroni post hoc test)

**Figure 5:**
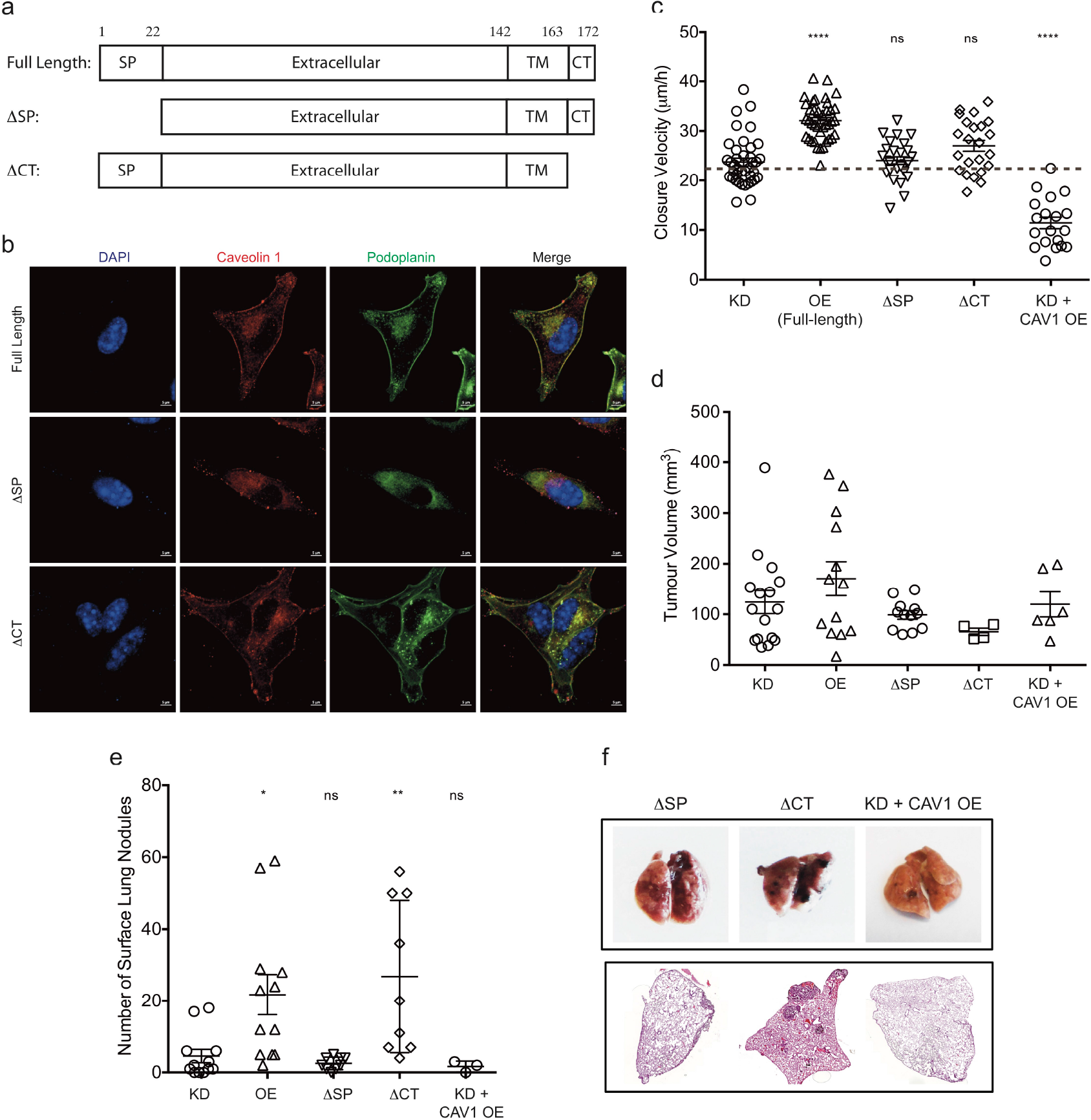
Podoplanin interacts with caveolin-1 through its transmembrane domain. a) Schematic of podoplanin variants: Full-length protein; ΔSP, protein without the N-terminal signal peptide; ΔCT, protein without the C-terminal cytoplasmic domain. SP, signal peptide (aa 1-22); Extracellular, extracellular domain (aa 23-142); TM, transmembrane domain (aa 143-163); CT, cytoplasmic domain (aa 164-172). b) Representative confocal images of podoplanin (green) and caveolin-1 (red) localization in the podoplanin variants. Nuclei counterstained with DAPI (blue). Scale bar: 5μm. c) Quantification of directional migration after wounding. Data presented as linear closure velocity of the scratch. n=3 independent experiments performed in triplicate. d) Primary tumour volumes 9 days after subcutaneous implantation of podoplanin-variants. n= at least 2 independent experiments. e) Quantification of tumour burden from lungs with metastatic nodules following intravenous injection of B16.F10 variants. n= at least 2 independent experiments. f) Representative images of tumour-bearing lungs 21 days after intravenous injection (top), and heamatoxylin/eosin stain images of lung sections illustrating tumour burden (bottom). Data presented as Mean ± SEM. *p<0.05, **p<0.01, ****p<0.0001, ns p>0.05 (One-way ANOVA with Bonferroni post hoc test). For c) data was compared to KD.

### Podoplanin transmembrane domain mediates caveolin-1 interactions

To examine the interactions between podoplanin and caveolin-1 in greater detail, and downstream consequences, further B16.F10 variants were generated in which endogenous podoplanin was knocked down and reconstituted with mutant forms (Fig. 5a and Fig. S3). We created two variants either lacking the short cytoplasmic domain, removing intracellular signalling capacity, or lacking the signal peptide, preventing podoplanin trafficking to the cell surface. A third variant, in which podoplanin deficient cells were transfected with caveolin-1 was also generated to identify podoplanin-independent but caveolin-1 dependent effects. Confocal analysis confirmed membrane co-localization of podoplanin with caveolin-1 (Fig. 5b top panel). Moreover, it confirmed that in the absence of signal peptide (ΔSP), podoplanin failed to reach the cell membrane, remaining cytoplasmic. Notably, in the absence of surface podoplanin, the majority of caveolin-1 also remained intracellular (Fig. 5b middle panel). In contrast, without the cytoplasmic domain, podoplanin again reached the cell surface co-localizing with caveolin-1 (Fig. 5b lower panel). As previously demonstrated, wound closure velocity increased with the level of cell surface podoplanin (Fig. 5c). In ΔSP cells, closure velocity of B16.F10 cells was significantly impaired, returning to levels comparable with knockdown variants. Similarly, variants expressing surface, but signalling-incompetent podoplanin (ΔCT) exhibited significantly retarded migration velocities, even with surface caveolin-1. Reconstitution with caveolin-1 in the absence of podoplanin was not capable of restoring migration further confirming that caveolin-1 was not the dominant driver. In fact, velocities were further reduced indicative of a dual function for caveolin-1 which in this case, may be tumour suppressive in the absence of podoplanin. These data point to the requirement of surface expression of signalling competent podoplanin for directional tumour migration, which is dependent on interaction and functional cooperation with caveolin-1. This interaction is likely to occur via the transmembrane domain. *In vivo*, the type of podoplanin (functional status or cellular location) had no impact on primary tumour growth (Fig. 5d), but the extracellular domain of podoplanin was sufficient for colonization (Fig. 5e) since both KD+CAV1 OE and ΔSP exhibited highly inefficient colony formation (Fig. 5e). In contrast, cells lacking the cytoplasmic tail formed lung nodules comparable to signalling competent podoplanin (Fig. 5e). These findings indicate that functional cooperation between podoplanin and caveolin-1 assists egress from the primary tumour through directional migration and invasion rather than a controlling growth or secondary colonization. Here, the formation of emboli through cell-cell or cell-platelet interactions may be sufficient for arrest in lungs and subsequent nodule formation.

### Cytoplasmic signalling potential of podoplanin

To address the question as to whether podoplanin transduces signals from outside to inside, we then focused on its small cytoplasmic domain. Previous studies indicated that podoplanin interacts with the cytoskeleton via the ERM (Ezrin, Radixin, Moesin) protein family and it is this path that controls migration (Martin-Villar et al., 2006). However, other reports described that with only 10 amino acids, the cytoplasmic domain may signal via phosphorylation of two serines by PKA and CDK5 (Krishnan et al., 2013; Krishnan et al., 2015). CDK5 (Cyclin-dependent kinase 5) is a cytoskeleton regulator reported to support melanoma cell motility and invasiveness (Bisht et al., 2015). In our cells, CDK5 is deregulated, increasing with podoplanin (Fig. 6a), yet intriguingly, siRNA disruption enhanced motility (Fig. 6b). While CDK5 levels increased at the protein level with podoplanin (Fig. 6a) this was not mirrored at the mRNA level (data not shown). This inverse relation to migration capacity may be explained by recently described functions of CDK5, phosphorylation of S171 of podoplanin inhibited migration (Krishnan et al., 2015). Thus in B16.F10 cells, accumulation beyond basal levels required for podoplanin phosphorylation may have a non-migration related function. To further evaluate the requirement for phosphorylation of podoplanin in execution of directional migration, we created podoplanin phospho-variants, transforming serine 167 into non-phosphorylatable (S167A) or constitutively active phosphomimetic (S167E) forms. In both phospho variants, serine 171 remained phosphorylatable. Migration in response to physical stimulation was impaired in S167A variants, resembling podoplanin deficient cells (Fig. 6c). Reconstitution with S167E had an even more profound impact on directional migration (Fig. 6c-d). Together, these data highlight that the cytoplasmic domain can indeed signal directly to the cytoskeleton to modulate migratory responses via CDK5 dependent phosphorylation. Finally, we investigated interactions between podoplanin variants and ERM proteins. As expected, increasing podoplanin levels corresponded to increased interactions with ERM proteins, which didn’t happen in ΔCT variant due to lack of its cytoplasmic domain (Fig. 6e). Interestingly, no differences were observed with phospho-serine variants indicating that ERM signalling is independent of phosphorylation status, instead relying on KKXXXR motifs.

**Figure 6:**
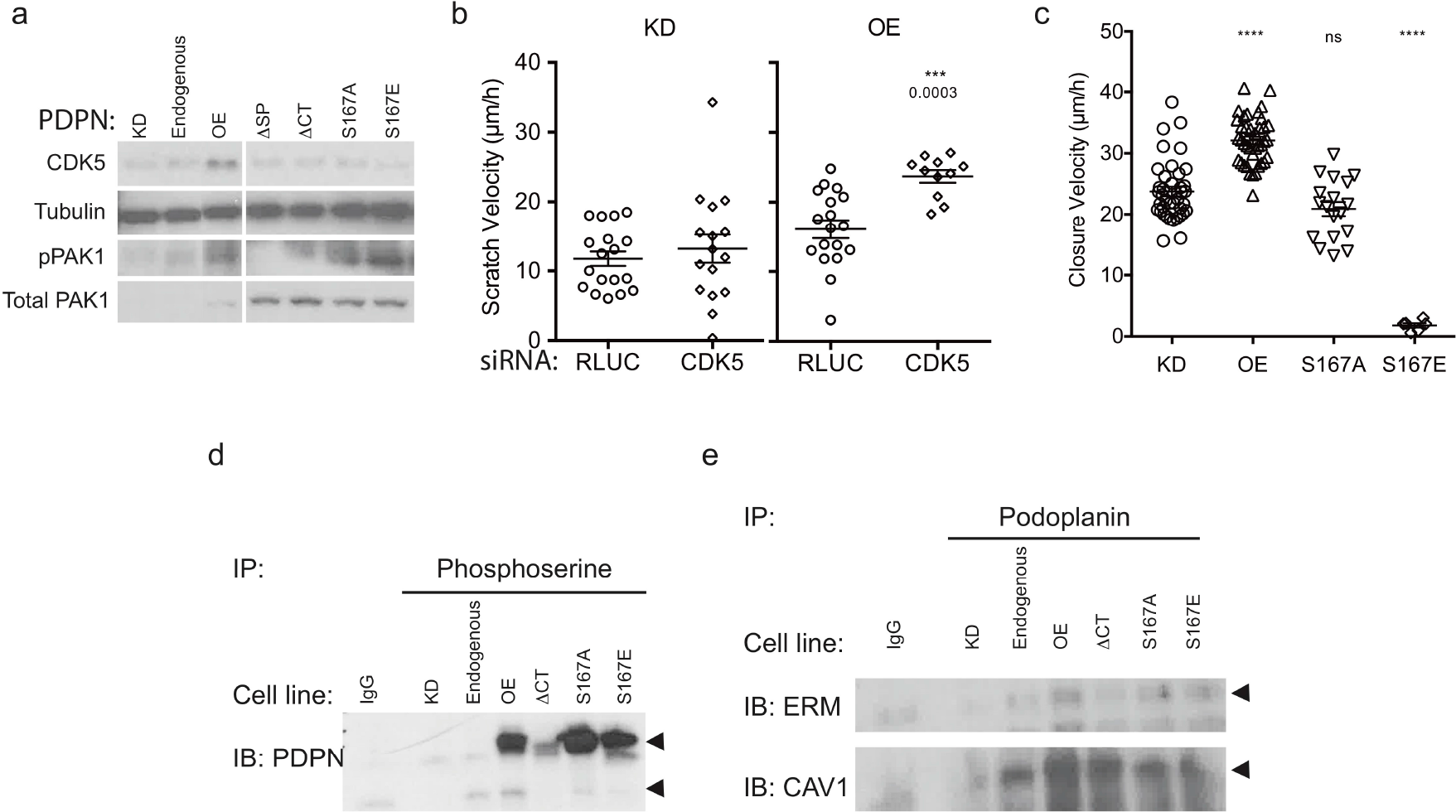
Podoplanin localization and phosphorylation status is a determinant of the migration capacity. a) Representative western blot detection of pPAK1, Total PAK1, CDK5, and Tubulin for all B16.F10 variants. b) Quantification of directional migration following wounding. Data presented as linear scratch closure velocity for cells KD and OE cells silenced with siRNA for CDK5. Data from 3 independent experiments performed in duplicate. c) Directional migration of KD, OE and phospho variants; S167A, phospho-mutant; S167E, phospho-mimetic. Data presented as linear scratch closure velocity. Data from 3 independent experiments performed in triplicate. d) Immunoprecipitation of phospho-serines to determine the degree of phosphorylation of podoplanin for the different podoplanin level and variant expressing cells. e) Immunoprecipitation analysis of the different podoplanin variants to determine interaction with the ERM protein family and caveolin-1. Data presented as Mean ± SEM. **p<0.001(Student t-test) (b), ****p<0.0001, ns p>0.05 (One-way ANOVA with Bonferroni post hoc test) (c).

We have demonstrated that podoplanin is able to influence tumour cell migration via a combination of elements: novel interactions with caveolin-1 or with ERM complexes, which transduce signals from the surface to the cytoskeleton. This is mainly accomplished through a balance of phosphorylation but also via PAK1-ERK1/2-ERM pathways, independent of small Rho GTPases activity.

## Discussion

Recent reports have demonstrated that expression of podoplanin expression in tumours correlates with poor prognosis (Martin-Villar et al., 2005; Ochoa-Alvarez et al., 2012; Wicki et al., 2006). Podoplanin induced RhoA activation and epithelial-mesenchymal transition (EMT) in MDCK canine cells (Martin-Villar et al., 2006), yet in MCF7 breast carcinoma cells, it attenuated RhoA activation, and although promoting invasion this was not via EMT (Wicki et al., 2006). Podoplanin is detected in numerous cell types including lymphatic endothelial cells, alveolar epithelial cells, fibroblastic reticular cells, podocytes and osteoblasts. Given the diversity of expression patterns and contradictory data regarding function, it is likely that podoplanin function is cell and context dependent. Moreover, many studies examining podoplanin function to date have been performed in cell lines lacking endogenous podoplanin expression (Martin-Villar et al., 2006; Wicki et al., 2006). Hence we investigated function in the context of behaviours that promote tumour progression using B16.F10 melanoma cells that endogenously express podoplanin. Podoplanin exerted no growth advantage to primary tumours either *in vitro* or *in vivo*, yet increasing levels supported significantly higher degrees of lung colonization consistent with previous studies in transfected CHO cells (Kunita et al., 2007) and MCF7 cells, which exhibited enhanced metastasis in the absence of accelerated growth. Here, podoplanin supported tumour lymphangiogenesis and metastasis to lymph nodes (Cueni et al., 2010). Superior capacity to colonize sites such as the lung is now understood to be mediated through platelets via interactions with the C-type lectin CLEC-2 (Kato et al., 2008; Suzuki-Inoue et al., 2007), which convey protective effects through release of growth factors to sustain tumour cell growth, providing physical support for cell immobilization, shielding them from high shear stresses, and limiting anti-tumour cell immune surveillance mechanisms (Labelle and Hynes, 2012; Palumbo et al., 2005). Here, podoplanin has a function in promoting metastasis due to its interaction with the extracellular environment. Our data support the view that it is the physical characteristics occurring as a consequence of surface podoplanin (i.e. platelet-tumour emboli formation) rather than any active signalling processes that underlie lung colonization since signalling incompetent variants still efficiently formed nodules.

Podoplanin had no influence on proliferation of B16.F10 cells in contrast to cells from different tissues and sites (Astarita et al., 2015), which in B16.F10 cells is driven by alternative signalling pathways including activation of FAK and Akt (Goundiam et al., 2010). In agreement with other studies, podoplanin influenced migration and invasion of cells which express it. However, this was only true upon receipt of a physical stimulus, having no impact on random migration of cells in a resting state. This implies that podoplanin has the potential to act as a microenvironment sensor, translating physical cues into an actionable, migration phenotype. To migrate in response to a stimulus cells require cell polarization, the formation of membrane extensions such as lamellipodia and filopodia, translocation of the cell body, and efficient mechanisms of release at the rear of the cell. These processes are tightly regulated by spatial activation of small GTPases Rho, Rac and Cdc42. Podoplanin expression is concentrated in actin-rich microvilli and plasma membrane projections including filopodia, lamellipodia and ruffles indicating a direct role in migration phenotypes (Li et al., 2015; Navarro et al., 2008; Scholl et al., 1999; Wicki et al., 2006) and has been further correlated with cytoskeleton and extracellular matrix remodelling to induce motility of individual cells (Cueni et al., 2010; Scholl et al., 1999). Although podoplanin has been implicated in collective cell migration (Cueni et al., 2010; Wicki et al., 2006) we observed expression to induce a more mesenchymal mode of migration. While podoplanin-induced migration has been proposed to occur via recruitment of ERM proteins and downstream activation of RhoA to drive EMT-like behaviour (Acton et al., 2014; Martin-Villar et al., 2006; Scholl et al., 1999), or via the inactivation or down-regulation of RhoA, Rac1 and Cdc42, to support collective migration (Cueni et al., 2010; Wicki et al., 2006), in endogenously expressing cells the migration phenotype was independent of these small GTPases. Instead, PAK1 emerged as the driver of podoplanin-induced migration in B16.F10 melanoma. Of note, PAK1 has been shown to drive increased invasion potential through cytoplasmic reorganization via the MAPK pathway in melanoma (Ong et al., 2013) and via F-actin rearrangements (Manser et al., 1997; Staser et al., 2013) or regulation of focal adhesion dynamics in other cell types (Arias-Romero and Chernoff, 2008; Slack-Davis et al., 2003). Together, these data imply that that podoplanin may be sufficient to act as an activator of PAK1 and therefore induce migration in tumour cells via the PAK1-ERM axis, bypassing the need for Rac1 activation.

Beyond colonization of secondary sites via platelet activation and aggregation (Kato et al., 2008; Suzuki-Inoue et al., 2007), it has been reported that pro-tumour functions of podoplanin arise following interactions with other membrane proteins such as CD44, which assisted with tethering tumour cells to ECM components (Martin-Villar et al., 2010; Tsuneki et al., 2013). In melanoma cells, podoplanin-induced migration in response to stimulus relied on its surface localization (Fernandez-Munoz et al., 2011) and physical interactions with caveolin-1 within lipid rafts, unlike in embryonic mouse alveolar epithelial cells (Barth et al., 2010). These opposing observations further highlight the context dependent nature of podoplanin behaviour. The interaction between podoplanin and caveolin-1 was fundamental to the migration capacity and phenotype of cells examined. Since both components have potential roles in formation and regulation of invadopodia, particularly relating to traffic of its constituents such as matrix metalloproteinases (Martin-Villar et al., 2015; Yamaguchi et al., 2009) and invadopodia stability (Martin-Villar et al., 2015), invadopodia turnover may be one mechanism by which the interaction is transduced. Indeed, we observed increased activity of matrix metalloproteinases in podoplanin expressing cells (Table 1 and data not shown).

Although caveolin-1 is key to vesicular transport, endocytosis and signal transduction (Martinez-Outschoorn et al., 2015), more recent pro-tumour activities including invadopodia formation, proliferation, migration and tumour forming capacity (Felicetti et al., 2009) have been reported following its activation (Lobos-Gonzalez et al., 2013; Senetta et al., 2013). As with podoplanin, its expression can be found localized to the leading edge are exposed to mechanical cues, such as stresses associated with adjacent stroma (Felicetti et al., 2009; Yoo et al., 2003). We demonstrated that directional migration responses required both caveolin-1 and podoplanin to be present at the surface since disruption to either significantly impaired the response to stimulation. However, although essential, restoration of caveolin-1 in the absence of podoplanin was not sufficient to rescue migratory phenotypes implying that podoplanin is the dominant partner driving these responses, likely directing caveolin-1 to its correct cell surface locale (Fig 4d) where any pro-tumour functions can occur (Felicetti et al., 2009; Yamaguchi et al., 2009).

Although surface expression of the podoplanin extracellular domain was sufficient for lung colonization, directional migration required both cell surface podoplanin and an intact cytoplasmic tail, indicating a requirement for signal transduction following stimulation. Within the 10 amino acid cytoplasmic domain, two potential serine phosphorylation sites (S167 and S171) exist. Their phosphorylation and subsequent downstream signalling has been described to be mediated through cooperation between PKA and CDK5 (Krishnan et al., 2013; Krishnan et al., 2015) whereby the default resting state of podoplanin is a phosphorylated to suppress migration, and de-phosphorylation results in promotion of a migration phenotype (Krishnan et al., 2013; Krishnan et al., 2015). Hence an increased migration of tumour cells may depend on the balance between the total amounts of podoplanin being expressed, but also the degree of phosphorylation and downstream substrates. In the case of B16.F10, we not only have an increase in podoplanin expression but also an increase in the expression in CDK5; yet we do not see a reduction in motility in response to stimulation as might be expected from previous studies in transfected cells (FIG 6a). While siRNA experiments where blockage of CDK5, substrate for S167, enhanced migration ability consistent with previous studies (Krishnan et al., 2013; Krishnan et al., 2015), mutation to a non-phosphorylatable form surprisingly impaired migration, though not to levels observed with constitutively phosphorylated variants (FIG 6c). Although we were not able to ascertain the phosphorylation status of S171 or PKA levels, the data implies the existence of alternate pro-migratory roles for CDK5 in these cells, so how may the CDK5 function in these cells beyond S167 phosphorylation? CDK5 is a proline-directed serine/threonine-protein kinase that has also been related to motility, invasiveness and metastatic spread in melanoma. It has been suggested that plays important roles in anchorage-independent growth, cell morphology, and in the phosphorylation of proteins required for actin reorganization, endocytosis and exocytosis (Bisht et al., 2015; Krishnan et al., 2015). In B16.F10, CDK5 may also directly modify downstream effectors of cytoskeleton, adhesion and migration including PAK1 (Strock et al., 2006) and ERK1 to modulate migration rather than inactivating podoplanin by serine phosphorylation. Here, we demonstrated that podoplanin may act through PAK1 and CDK5 to promote cell migration whereby CDK5 is sufficient to activate PAK1, independent of the Rho-GTPases, and therefore interact with the cytoskeleton via the ERM protein family. Furthermore, podoplanin-induced migration supported by increased matrix degradation was accomplished only the presence of caveolin-1 at the cell surface. Since invadopodia formation is regulated by caveolin-1–mediated control of membrane cholesterol levels (Caldieri et al., 2009; Yamaguchi et al., 2009), and podoplanin has been identified within invadopodia adhesion rings critical to proteolytic degradation of the extracellular matrix (Martin-Villar et al., 2015; Yamaguchi et al., 2009), our data are consistent with the idea that podoplanin and caveolin-1 cooperation relates to invadopodia turnover (Martin-Villar et al., 2015; Yamaguchi et al., 2009) to drive directional motility responses.

In summary, we have demonstrated that functional cooperation between cell surface podoplanin and caveolin-1 are required for directional migration and invasion of melanoma cells via CDK5, PAK1 and ERK1 mediated signal transduction.

## Materials and Methods

### Cell culture

B16.F10 mouse melanoma cells (ATCC) were cultured in DMEM supplemented with 10U/mL of penicillin, 10μg/mL of streptomycin, and 10% FBS, maintained in a humidified 5% CO2 atmosphere incubator at 37°C. Viability were assessed by trypan blue dye-exclusion.

### Cloning of podoplanin variants and caveolin-1 plasmid

Insert amplifications (Table S1-S2) were performed using Phusion DNA Polymerase (Thermo Scientific) according to manufacturer instructions. Inserts were digested with FastDigest restriction enzymes MluI and SfiI (Fermentas). Ligation to pCMV6-AC-tGFP (Origene) was performed using T4 DNA ligase (New England Biolabs). NEB10-beta competent *E.coli* cells (New England Biolabs) were transformed by heat shock and selected with kanamycin. Positive colonies were confirmed by sequencing.

### Transfections

Cells were transfected with Fugene6 (Promega), as per manufacturer instructions. Silencing was performed using podoplanin-targeted and control shRNA lentiviral transduction particles (Sigma Aldrich, MISSION^®^ shRNA Lentiviral Transduction Particles, ref. SHCLNV-NM_010329 (TRCN0000174621) and SHC002V, respectively) at a MOI of 1 and selected with puromycin. Resistant cells were transfected with podoplanin variant constructs and selected for G418 resistance. Transient knockdowns were achieved by reverse-transfecting cells with esiRNA (Sigma) against PDPN (cat# EMU061611), CAV1 (cat# EMU013051), CDK5 (cat# EMU063451), PAK1 (cat# EMU026031) and control siRNA RLUC (cat# EHURLUC). Transfections were performed using Lipofectamine RNAiMax reagent (Life Technologies) as per manufacturer instructions.

### Expression profiling

RT^2^ First Strand kit (Qiagen) for cDNA synthesis was used to reverse transcribe 1μg of total RNA. PCR arrays targeting mouse cell motility (ref. PAMM-128) and cytoskeleton regulators (ref. PAMM-088; SABiosciences) were performed as per manufacturers guidelines. Quantitative RT-PCR was performed using RT^2^ SYBR Green Mastermix and run on and LC480 cycler (Roche). Post-analysis was carried out using the RT^2^ Profiler PCR Data Online Analysis tool (SABiosciences).

### Random migration assay

Cells were seeded in chamber slides. After attaching overnight, brightfield live imagining of randomly selected fields (100x total magnification) was performed using a Live Cell imaging microscope (AF7000, Leica Biosystems) for 24 hours. Migration analysis was performed using Volocity Software (Perkin Elmer).

### Wound healing scratch assay

Cells were seeded in an 8-well chamber and left to attach overnight. A scratch was then performed with the aid of a pipette tip. Sequential brightfield images of each scratch were taken by a live-cell imaging microscope (AF7000, Leica Biosystems) for 24 hours in randomly selected fields (100x total magnification). Migration analysis and closure velocity were performed using Volocity Software (Perkin Elmer).

### Spheroid invasion assay

Spheroids of 500 cells were generated via the methylcellulose hanging drop method (Ware et al., 2016) 24 hours prior to seeding them into 2mg/mL type I rat tail collagen gels (BD Biosciences) on borosilicate chamber slides. Live imaging of randomly selected spheroids was performed over 72 hours using a Live Cell imaging microscope (AF7000, Leica Biosystems). Post-analysis was performed using Volocity Software (Perkin Elmer).

### Flow cytometry

Cells were detached using Trypsin-EDTA (Sigma), pelleted and re-suspended in PBS with 0.5%(w/v) BSA. For surface marker staining, cells were incubated with PDPN-APC (1:400, Biolegend) for 30 minutes on ice. Cells were washed twice before characterization (Fortessa, BD) or sorting (BD). Results were analysed using FlowJo X 10.0.7r2 software (Treestar).

### Proliferation and viability measurement

#### BrdU measurement

Cell proliferation was analysed using a eBioscience BrdU staining kit for flow cytometry and performed according to manufacturer instructions. Cells used for the assay were cultured overnight in a 6-well plate. Data was acquired using a Fortessa flow cytometer (BD) and analyzed offline with Flowjo software (Treestar). *MTT cell proliferation assay:* Cells were cultured overnight in 24-well plates. On the day of the assay cells were washed with PBS and then incubated with 0.45 mg/mL of Thiazolyl Blue Tetrazolium Bromide (Sigma) in PBS. Cells were then incubated for 4 hours. Formazan crystals were then dissolved in DMSO and quantified by absorbance reads at 570 and 700 nm in a multi-plate reader (Tecan). *Cell Titer Blue cell viability assay:* Cell viability was determined according to manufacturer instructions (Promega), in a 24-well plate. Fluorescence at 560/590 nm was measured in a multi-plate reader (Tecan).

### Active protein pull-down and detection

Rho, Rac1 and Cdc42 active protein measurement was performed using Pierce pull-down kits (Thermo Scientific, Cat. numbers 16116, 16118, and 16119, respectively), as per manufacturer instructions. The presence of the active protein was determined by western blot of elutes (see Table S3 for conditions) in cells that were stimulated with a scratch.

### Immunoprecipitation

Adherent cells were washed with cold PBS twice then scraped directly into cold lysis buffer (20mM Tris-HCl pH 7.4, 0.2% NP-40, 150mM NaCl, 2mM sodium orthovanadate, and protease inhibitors cocktail (Roche)). 1μg of the pull-down antibody (Podoplanin or Caveolin-1) was added to 1.5 mg of total protein incubated for 3 hours at 4°C. Protein G Plus Agarose beads (Santa Cruz Biotechnology) were added before incubation for a further hour. The precipitated conjugates attached to beads were washed three times with lysis buffer containing 1% Triton X-100. Precipitates were then removed from the beads by incubation with protein loading buffer (50mM Tris-HCl pH6.8, 2% SDS, 10% glycerol, 1% beta-mercaptoethanol, 12.5mM EDTA, 0.02% bromophenol blue) for 5 minutes and boiling for 10 minutes. Samples were analysed by western blot (Table S3).

### Membrane raft extraction

Lipid raft extraction was performed by gradient ultracentrifugation as previously described by George et al. (George et al., 2010).

### Immunoblotting

Protein samples were separated by 12% SDS-PAGE and transferred onto a PVDF membrane (Millipore). The membranes were blocked with 5% BSA in TBS containing 0.5% Tween 20 and then incubated with primary antibodies overnight at 4°C (Table S3). Washed membranes were incubated with the appropriate HRP-conjugated secondary antibody prior to detection with an enhanced chemiluminescence detection kit (Pierce).

### In vivo studies

Experiments involving animals were performed in accordance with UK Home Office regulations under HO license PPL 80/2574. Where possible, experimental groups were randomized and blinded. Lung colonization capacity was determined by intravenous injection of 150,000 B16.F10 cells into 7-week old female C57BL/6 mice (Envigo). Animals were sacrificed 21 days later and lungs analysed for nodules and tumour burden. Samples were processed for histology. For primary tumour studies, 250,000 cells were implanted subcutaneously on both shoulders of female C57BL/6 mice and growth was monitored for 9 days or until tumour sizes reached the legal limit of a diameter of 12 mm. Tumour volumes were calculated by: Volume=π/6×small^2^×longest.

### Haematoxylin and eosin stain

Harvested lungs were embedded in OCT medium (TissueTek). 10μm sections were fixed in 4% para-formaldehyde. Hematoxylin and eosin stain was performed using a Leica Autostainer XL. Slides were mounted with a Leica CV5030 and imaged using AxioScan.Z1 (Zeiss) slide scanner. Tumour burden was determined using Fiji software. Percentage of tumour burden calculated as tumour area/total area × 100.

### Immunofluorescence

Cells were fixed in cold methanol, blocked with 5% chicken serum in PBS, and incubated with the appropriate primary antibodies overnight (Table S3). Conjugated secondary antibodies were incubated at room temperature. Nuclei were counterstained with DAPI and slides were mounted in SlowFade Gold antifade reagent (Life Technologies). Confocal images were taken using either Leica SP5 or Zeiss 880 and processed with Volocity (Perkin Elmer) or Zen Software (Zeiss).

### PAK1 chemical inhibition

Cells were allowed to attach prior to treatment for 48 hours with 2.5, 5, and 10μM IPA-3 (Sigma). Treated cells were assessed for proliferation and viability, imaged by confocal microscopy, and scratched for directional migration assessment.

### Statistical analysis

Statistical analyses were performed using GraphPad Prism 6 software (GraphPad). For comparisons of two groups, Students t-tests were performed. When comparing 3 or more datasets, One-way ANOVA and appropriate post-hoc tests were performed.

## Supporting information

## Acknowledgements

The authors would like to thank ARES facility staff for assistance with *in vivo* experiments, and members of the CIMR flow cytometry core for assistance with cell sorting applications.

## References

Acton, S.E., Astarita, J.L., Malhotra, D., Lukacs-Kornek, V., Franz, B., Hess, P.R., Jakus, Z., Kuligowski, M., Fletcher, A.L., Elpek, K.G., et al. (2012). Podoplanin-rich stromal networks induce dendritic cell motility via activation of the C-type lectin receptor CLEC-2. Immunity 37, 276–289.

Acton, S.E., Farrugia, A.J., Astarita, J.L., Mourao-Sa, D., Jenkins, R.P., Nye, E., Hooper, S., van Blijswijk, J., Rogers, N.C., Snelgrove, K.J., et al. (2014). Dendritic cells control fibroblastic reticular network tension and lymph node expansion. Nature 514, 498–502.

Arias-Romero, L.E., and Chernoff, J. (2008). A tale of two Paks. Biol Cell 100, 97–108.

Astarita, J.L., Cremasco, V., Fu, J., Darnell, M.C., Peck, J.R., Nieves-Bonilla, J.M., Song, K., Kondo, Y., Woodruff, M.C., Gogineni, A., et al. (2015). The CLEC-2-podoplanin axis controls the contractility of fibroblastic reticular cells and lymph node microarchitecture. Nat Immunol 16, 75–84.

Barth, K., Blasche, R., and Kasper, M. (2010). T1alpha/podoplanin shows raft-associated distribution in mouse lung alveolar epithelial E10 cells. Cell Physiol Biochem 25, 103–112.

Bisht, S., Nolting, J., Schutte, U., Haarmann, J., Jain, P., Shah, D., Brossart, P., Flaherty, P., and Feldmann, G. (2015). Cyclin-Dependent Kinase 5 (CDK5) Controls Melanoma Cell Motility, Invasiveness, and Metastatic Spread-Identification of a Promising Novel therapeutic target. Transl Oncol 8, 295–307.

Caldieri, G., Giacchetti, G., Beznoussenko, G., Attanasio, F., Ayala, I., and Buccione, R. (2009). Invadopodia biogenesis is regulated by caveolin-mediated modulation of membrane cholesterol levels. J Cell Mol Med 13, 1728–1740.

Cueni, L.N., Hegyi, I., Shin, J.W., Albinger-Hegyi, A., Gruber, S., Kunstfeld, R., Moch, H., and Detmar, M. (2010). Tumor Lymphangiogenesis and Metastasis to Lymph Nodes Induced by Cancer Cell Expression of Podoplanin. American Journal of Pathology 177, 1004–1016.

Felicetti, F., Parolini, I., Bottero, L., Fecchi, K., Errico, M.C., Raggi, C., Biffoni, M., Spadaro, F., Lisanti, M.P., Sargiacomo, M., et al. (2009). Caveolin-1 tumor-promoting role in human melanoma. Int J Cancer 125, 1514–1522.

Fernandez-Munoz, B., Yurrita, M.M., Martin-Villar, E., Carrasco-Ramirez, P., Megias, D., Renart, J., and Quintanilla, M. (2011). The transmembrane domain of podoplanin is required for its association with lipid rafts and the induction of epithelial-mesenchymal transition. Int J Biochem Cell Biol 43, 886–896.

George, K.S., Wu, Q., and Wu, S. (2010). Effects of freezing and protein cross-linker on isolating membrane raft-associated proteins. Biotechniques 49, 837–838.

Goundiam, O., Nagel, M.D., and Vayssade, M. (2010). Growth and survival signalling in B16F10 melanoma cells in 3D culture. Cell Biol Int 34, 385–391.

Herzog, B.H., Fu, J., Wilson, S.J., Hess, P.R., Sen, A., McDaniel, J.M., Pan, Y., Sheng, M., Yago, T., Silasi-Mansat, R., et al. (2013). Podoplanin maintains high endothelial venule integrity by interacting with platelet CLEC-2. Nature 502, 105–109.

Kaneko, M.K., Kunita, A., Abe, S., Tsujimoto, Y., Fukayama, M., Goto, K., Sawa, Y., Nishioka, Y., and Kato, Y. (2012). Chimeric anti-podoplanin antibody suppresses tumor metastasis through neutralization and antibody-dependent cellular cytotoxicity. Cancer Sci 103, 1913–1919.

Kato, Y., Kaneko, M.K., Kunita, A., Ito, H., Kameyama, A., Ogasawara, S., Matsuura, N., Hasegawa, Y., Suzuki-Inoue, K., Inoue, O., et al. (2008). Molecular analysis of the pathophysiological binding of the platelet aggregation-inducing factor podoplanin to the C-type lectin-like receptor CLEC-2. Cancer Sci 99, 54–61.

Kato, Y., Kaneko, M.K., Kuno, A., Uchiyama, N., Amano, K., Chiba, Y., Hasegawa, Y., Hirabayashi, J., Narimatsu, H., Mishima, K., et al. (2006). Inhibition of tumor cell-induced platelet aggregation using a novel anti-podoplanin antibody reacting with its platelet-aggregation-stimulating domain. Biochem Biophys Res Commun 349, 1301–1307.

Krishnan, H., Ochoa-Alvarez, J.A., Shen, Y., Nevel, E., Lakshminarayanan, M., Williams, M.C., Ramirez, M.I., Miller, W.T., and Goldberg, G.S. (2013). Serines in the intracellular tail of podoplanin (PDPN) regulate cell motility. J Biol Chem 288, 12215–12221.

Krishnan, H., Retzbach, E.P., Ramirez, M.I., Liu, T., Li, H., Miller, W.T., and Goldberg, G.S. (2015). PKA and CDK5 can phosphorylate specific serines on the intracellular domain of podoplanin (PDPN) to inhibit cell motility. Exp Cell Res 335, 115–122.

Kunita, A., Kashima, T.G., Morishita, Y., Fukayama, M., Kato, Y., Tsuruo, T., and Fujita, N. (2007). The platelet aggregation-inducing factor aggrus/podoplanin promotes pulmonary metastasis. Am J Pathol 170, 1337–1347.

Labelle, M., and Hynes, R.O. (2012). The initial hours of metastasis: the importance of cooperative host-tumor cell interactions during hematogenous dissemination. Cancer Discov 2, 1091–1099.

Li, Y.Y., Zhou, C.X., and Gao, Y. (2015). Podoplanin promotes the invasion of oral squamous cell carcinoma in coordination with MT1-MMP and Rho GTPases. Am J Cancer Res 5, 514–529.

Lobos-Gonzalez, L., Aguilar, L., Diaz, J., Diaz, N., Urra, H., Torres, V.A., Silva, V., Fitzpatrick, C., Lladser, A., Hoek, K.S., et al. (2013). E-cadherin determines Caveolin-1 tumor suppression or metastasis enhancing function in melanoma cells. Pigment Cell Melanoma Res 26, 555–570.

Manser, E., Huang, H.Y., Loo, T.H., Chen, X.Q., Dong, J.M., Leung, T., and Lim, L. (1997). Expression of constitutively active alpha-PAK reveals effects of the kinase on actin and focal complexes. Mol Cell Biol 17, 1129–1143.

Martin-Villar, E., Borda-d'Agua, B., Carrasco-Ramirez, P., Renart, J., Parsons, M., Quintanilla, M., and Jones, G.E. (2015). Podoplanin mediates ECM degradation by squamous carcinoma cells through control of invadopodia stability. Oncogene 34, 4531–4544.

Martin-Villar, E., Fernandez-Munoz, B., Parsons, M., Yurrita, M.M., Megias, D., Perez-Gomez, E., Jones, G.E., and Quintanilla, M. (2010). Podoplanin associates with CD44 to promote directional cell migration. Mol Biol Cell 21, 4387–4399.

Martin-Villar, E., Megias, D., Castel, S., Yurrita, M.M., Vilaro, S., and Quintanilla, M. (2006). Podoplanin binds ERM proteins to activate RhoA and promote epithelial-mesenchymal transition. J Cell Sci 119, 4541–4553.

Martin-Villar, E., Scholl, F.G., Gamallo, C., Yurrita, M.M., Munoz-Guerra, M., Cruces, J., and Quintanilla, M. (2005). Characterization of human PA2.26 antigen (T1alpha-2, podoplanin), a small membrane mucin induced in oral squamous cell carcinomas. Int J Cancer 113, 899–910.

Martinez-Outschoorn, U.E., Sotgia, F., and Lisanti, M.P. (2015). Caveolae and signalling in cancer. Nat Rev Cancer 15, 225–237.

Navarro, A., Perez, R.E., Rezaiekhaligh, M., Mabry, S.M., and Ekekezie, I.I. (2008). T1 alpha/podoplanin is essential for capillary morphogenesis in lymphatic endothelial cells. Am J Physiol-Lung C 295, L543–L551.

Nose, K., Saito, H., and Kuroki, T. (1990). Isolation of a gene sequence induced later by tumor-promoting 12-O-tetradecanoylphorbol-13-acetate in mouse osteoblastic cells (MC3T3-E1) and expressed constitutively in ras-transformed cells. Cell Growth Differ 1, 511–518.

Ochoa-Alvarez, J.A., Krishnan, H., Shen, Y., Acharya, N.K., Han, M., McNulty, D.E., Hasegawa, H., Hyodo, T., Senga, T., Geng, J.G., et al. (2012). Plant lectin can target receptors containing sialic acid, exemplified by podoplanin, to inhibit transformed cell growth and migration. PLoS One 7, e41845.

Ong, C.C., Jubb, A.M., Jakubiak, D., Zhou, W., Rudolph, J., Haverty, P.M., Kowanetz, M., Yan, Y., Tremayne, J., Lisle, R., et al. (2013). P21-activated kinase 1 (PAK1) as a therapeutic target in BRAF wild-type melanoma. J Natl Cancer Inst 105, 606–607.

Palumbo, J.S., Talmage, K.E., Massari, J.V., La Jeunesse, C.M., Flick, M.J., Kombrinck, K.W., Jirouskova, M., and Degen, J.L. (2005). Platelets and fibrin(ogen) increase metastatic potential by impeding natural killer cell-mediated elimination of tumor cells. Blood 105, 178–185.

Prideaux, M., Loveridge, N., Pitsillides, A.A., and Farquharson, C. (2012). Extracellular matrix mineralization promotes E11/gp38 glycoprotein expression and drives osteocytic differentiation. PLoS One 7, e36786.

Scholl, F.G., Gamallo, C., Vilaro, S., and Quintanilla, M. (1999). Identification of PA2.26 antigen as a novel cell-surface mucin-type glycoprotein that induces plasma membrane extensions and increased motility in keratinocytes. J Cell Sci 112 (Pt 24), 4601–4613.

Senetta, R., Stella, G., Pozzi, E., Sturli, N., Massi, D., and Cassoni, P. (2013). Caveolin-1 as a promoter of tumour spreading: when, how, where and why. J Cell Mol Med 17, 325–336.

Shen, Y., Chen, C.S., Ichikawa, H., and Goldberg, G.S. (2010). SRC induces podoplanin expression to promote cell migration. J Biol Chem 285, 9649–9656.

Slack-Davis, J.K., Eblen, S.T., Zecevic, M., Boerner, S.A., Tarcsafalvi, A., Diaz, H.B., Marshall, M.S., Weber, M.J., Parsons, J.T., and Catling, A.D. (2003). PAK1 phosphorylation of MEK1 regulates fibronectin-stimulated MAPK activation. J Cell Biol 162, 281–291.

Staser, K., Shew, M.A., Michels, E.G., Mwanthi, M.M., Yang, F.C., Clapp, D.W., and Park, S.J. (2013). A Pak1-PP2A-ERM signaling axis mediates F-actin rearrangement and degranulation in mast cells. Exp Hematol 41, 56–66 e52.

Strock, C.J., Park, J.I., Nakakura, E.K., Bova, G.S., Isaacs, J.T., Ball, D.W., and Nelkin, B.D. (2006). Cyclin-dependent kinase 5 activity controls cell motility and metastatic potential of prostate cancer cells. Cancer Res 66, 7509–7515.

Suzuki, H., Kato, Y., Kaneko, M.K., Okita, Y., Narimatsu, H., and Kato, M. (2008). Induction of podoplanin by transforming growth factor-beta in human fibrosarcoma. FEBS Lett 582, 341–345.

Suzuki-Inoue, K., Kato, Y., Inoue, O., Kaneko, M.K., Mishima, K., Yatomi, Y., Yamazaki, Y., Narimatsu, H., and Ozaki, Y. (2007). Involvement of the snake toxin receptor CLEC-2, in podoplanin-mediated platelet activation, by cancer cells. J Biol Chem 282, 25993–26001.

Takagi, S., Sato, S., Oh-hara, T., Takami, M., Koike, S., Mishima, Y., Hatake, K., and Fujita, N. (2013). Platelets promote tumor growth and metastasis via direct interaction between Aggrus/podoplanin and CLEC-2. PLoS One 8, e73609.

Takakubo, Y., Oki, H., Naganuma, Y., Saski, K., Sasaki, A., Tamaki, Y., Suran, Y., Konta, T., and Takagi, M. (2016). Distribution of podoplanin in synovial tissues in rheumatoid arthritis patients using biologic or conventional disease-modifying anti-rheumatic drugs. Curr Rheumatol Rev.

Tsuneki, M., Yamazaki, M., Maruyama, S., Cheng, J., and Saku, T. (2013). Podoplanin-mediated cell adhesion through extracellular matrix in oral squamous cell carcinoma. Lab Invest 93, 921–932.

Uhrin, P., Zaujec, J., Breuss, J.M., Olcaydu, D., Chrenek, P., Stockinger, H., Fuertbauer, E., Moser, M., Haiko, P., Fassler, R., et al. (2010). Novel function for blood platelets and podoplanin in developmental separation of blood and lymphatic circulation. Blood 115, 3997–4005.

Ware, M.J., Colbert, K., Keshishian, V., Ho, J., Corr, S.J., Curley, S.A., and Godin, B. (2016). Generation of Homogenous Three-Dimensional Pancreatic Cancer Cell Spheroids Using an Improved Hanging Drop Technique. Tissue Eng Part C Methods 22, 312–321.

Wicki, A., Lehembre, F., Wick, N., Hantusch, B., Kerjaschki, D., and Christofori, G. (2006). Tumor invasion in the absence of epithelial-mesenchymal transition: podoplanin-mediated remodeling of the actin cytoskeleton. Cancer Cell 9, 261–272.

Yamaguchi, H., Takeo, Y., Yoshida, S., Kouchi, Z., Nakamura, Y., and Fukami, K. (2009). Lipid rafts and caveolin-1 are required for invadopodia formation and extracellular matrix degradation by human breast cancer cells. Cancer Res 69, 8594–8602.

Yoo, S.H., Park, Y.S., Kim, H.R., Sung, S.W., Kim, J.H., Shim, Y.S., Lee, S.D., Choi, Y.L., Kim, M.K., and Chung, D.H. (2003). Expression of caveolin-1 is associated with poor prognosis of patients with squamous cell carcinoma of the lung. Lung Cancer 42, 195–202.

