## Supplementary material for "Podoplanin interaction with caveolin-1 promotes tumour cell migration and invasion"

### Supplementary Tables

**Supplementary Table S1**: Primers for cloning podoplanin variants. Template: Plasmid pCMV6-mPdpn-tGFP (Origene, ref. MG201468)

| Pdpn Variant | Primer | Sequence | Tm (ºC) |
|---|---|---|---|
| Delta_SP | Fw | ATGGGGACTATAGGCGTGAATGAAGATGA TATTGT | 60.5 |
| | Rv | GGGCGAGAACCTTCCAGAAATCTTCTTC | 61.4 |
| Delta_CT | Fw | ATGTGGACCGTGCCAGTGTTGTTCTG | 61.1 |
| | Rv | AACAACAATGAAGATCCCTCCGACGAAGC | 61.5 |

**Supplementary Table S2**: Primers used for cloning of Caveolin-1. Template: cDNA created from mRNA extracted from B16.F10 cells

| | Primer | Sequence | Tm (ºC) |
|---|---|---|---|
| Cav1 | Fw | ATGTCTGGGGGCAAATACGTAGACTCC | 61.3 |
| | Rv | TATCTCTTTCTGCGTGCTGATGCGGATG | 61.4 |

**Supplementary Table S3**: Antibodies used for protein detection by western blot and immunofluorescence.

| Primary/Secondary antibody | Application | Dilution | Supplier |
|---|---|---|---|
| Anti-Podoplanin | WB | 1:1000 | R&D Systems (ref. BAF3244) |
| Anti-Podoplanin | Immunofluorescence | 1:100 | R&D Systems (ref. BAF3244) |
| Anti-Caveolin-1 | WB | 1:500 | Abcam (ref. 2910) |
| Anti-Caveolin-1 | Immunofluorescence | 1:100 | Abcam (ref. 2910) |
| Anti-Rho | WB | 1:500 | Pierce (ref. 1862332) |
| Anti-Rac 1 | WB | 1:500 | Pierce (ref. 1862341) |
| Anti-Cdc42 | WB | 1:500 | Pierce (ref. 1862345) |
| Anti-PAK1 | WB | 1:500 | Cell Signaling (ref. 2602) |
| Anti-pPAK1 | WB | 1:500 | Cell Signaling (ref. 9101) |
| Anti-pERK1/2 | WB | 1:500 | Cell Signaling (ref. 4370) |
| Anti-ERK1/2 | WB | 1:500 | Cell Signaling (ref. 9102) |
| Anti-Cdck5 | WB | 1:500 | GeneTex (ref. GTX108328) |
| Anti-Tubulin | WB | 1:2000 | Sigma (ref. T6074) |
| Anti-Goat HRP | WB | 1:1000 | DakoCytomaton (ref. P0160) |
| Anti-Rabbit | WB | 1:1000 | DakoCytomaton (ref. P0399) |
| Anti-Mouse | WB | 1:1000 | DakoCytomaton (ref. P0447) |

### Supplementary Figures

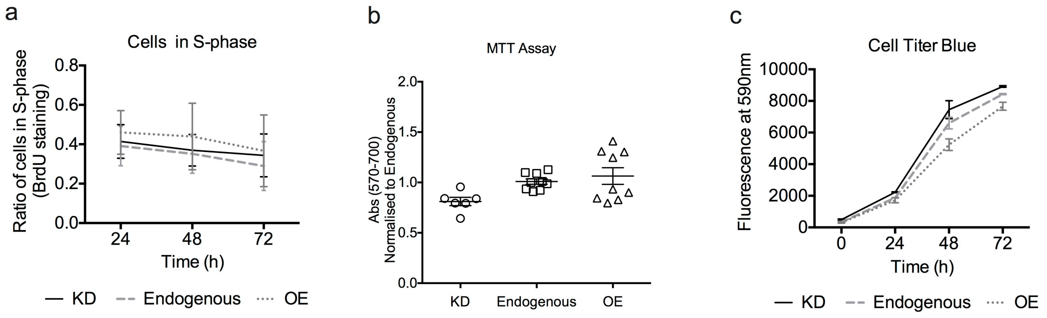


**Supplementary Figure S1**: a) BrdU proliferation assay for the cells with different podoplanin levels. Data from 2 independent experiments performed in triplicate. b) MTT metabolic assay for the cells with different podoplanin levels. Data from 3 independent experiments performed in triplicate. c) Cell titer blue viability assay for the cells with different podoplanin levels. Representative data from 3 independent experiments. Data presented as Mean ± SEM.

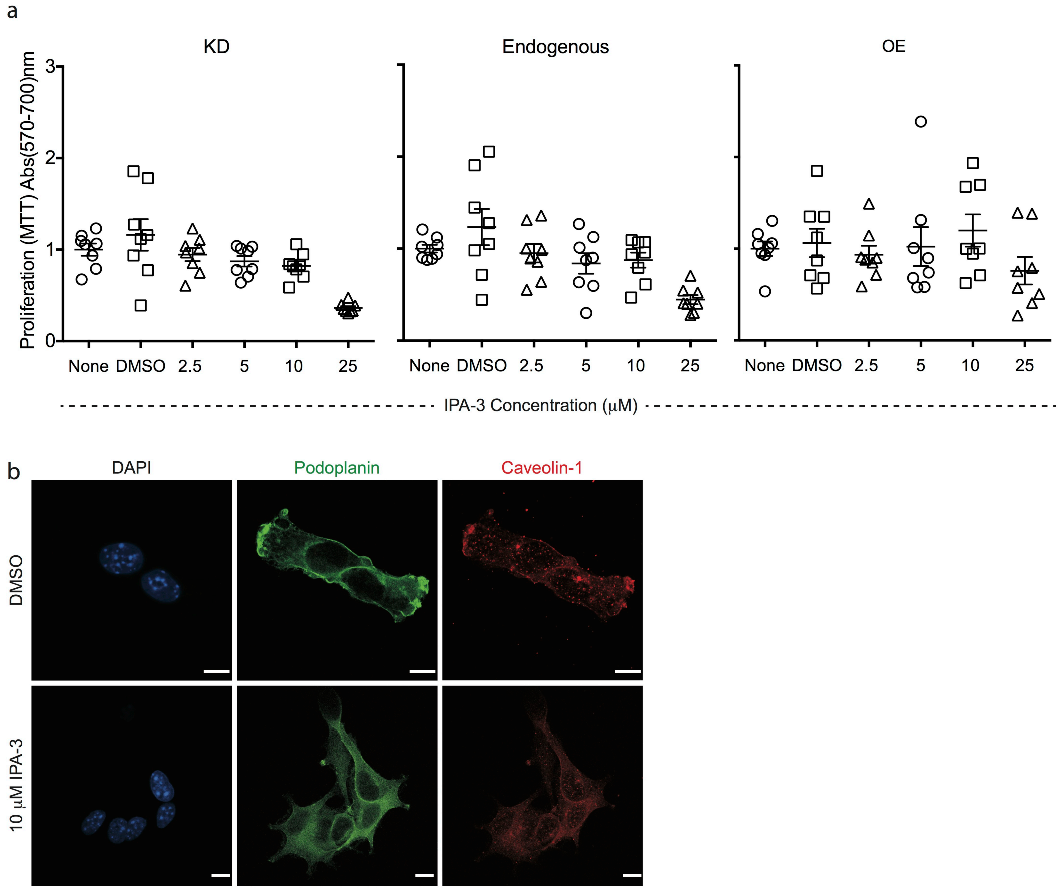


**Supplementary Figure S2**: a) Proliferation of cells with different levels of podoplanin levels in the presence of increasing concentrations of PAK1 inhibitor IPA-3. Higher concentrations of IPA-3 impact the proliferative ability of cells (also measured with AnnexinV staining – data not shown). b) Immunofluorescence for podoplanin (green) and caveolin-1 (red) of cells treated with 10 μm IPA-3 or DMSO control. Nuclei counterstained with DAPI (blue).

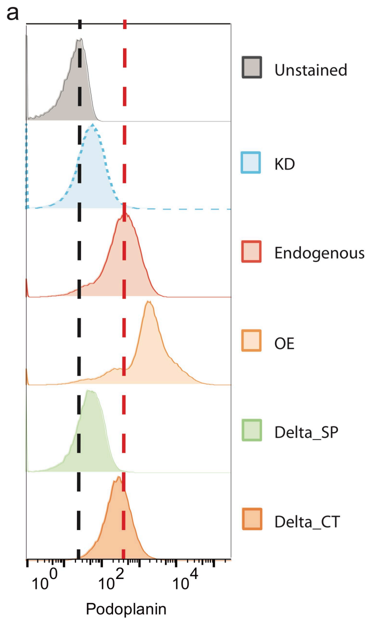


**Supplementary Figure S3:** a) Flow cytometry histograms illustrating surface podoplanin expression by the different podoplanin variant cell lines. Red dashed line indicates mean podoplanin levels in endogenous cells; Black line shows unstained control.
